# Puromycin reactivity does not accurately localize translation at the subcellular level

**DOI:** 10.1101/2020.06.22.165217

**Authors:** Syed Usman Enam, Boris Zinshteyn, Daniel H. Goldman, Madeline Cassani, Nathan M. Livingston, Geraldine Seydoux, Rachel Green

**Author notes:** These authors contributed equally (Syed Usman Enam, Boris Zinshteyn, Daniel H. Goldman). Correspondence to (R.G.).

## Abstract

Puromycin is a tyrosyl-tRNA mimic that blocks translation by labeling and releasing elongating polypeptide chains from translating ribosomes. Puromycin has been used in molecular biology research for decades as a translation inhibitor. The development of puromycin antibodies and derivatized puromycin analogs has enabled the quantification of active translation in bulk and single-cell assays. More recently, *in vivo* puromycylation assays have become popular tools for localizing translating ribosomes in cells. These assays often use elongation inhibitors to purportedly inhibit the release of puromycin-labeled nascent peptides from ribosomes. Here, using *in vitro* and *in vivo* experiments, we demonstrate that, even in the presence of elongation inhibitors, puromycylated peptides are released and diffuse away from ribosomes. Puromycylation assays reveal subcellular sites, such as nuclei, where puromycylated peptides accumulate post-release and which do not necessarily coincide with sites of active translation. Our findings urge caution when interpreting puromycylation assays in the *in vivo* context.

## Introduction

Puromycin is a potent translational inhibitor that binds to ribosomes from all domains of life and has been used as a chemical probe and selectable marker for decades (*1, 2*). Puromycin is unique among translational inhibitors in that it is itself a substrate of the ribosomal peptidyl-transferase reaction (*3*). Puromycin mimics the 3′ adenosine of a tRNA charged with a modified tyrosine, which binds in the ribosomal acceptor site (Figure 1A). The ribosome transfers the nascent peptide chain on the P-site tRNA to puromycin, leading to spontaneous dissociation of the nascent peptide from the ribosome (Figure 1B) (*3*).

**Fig. 1:**
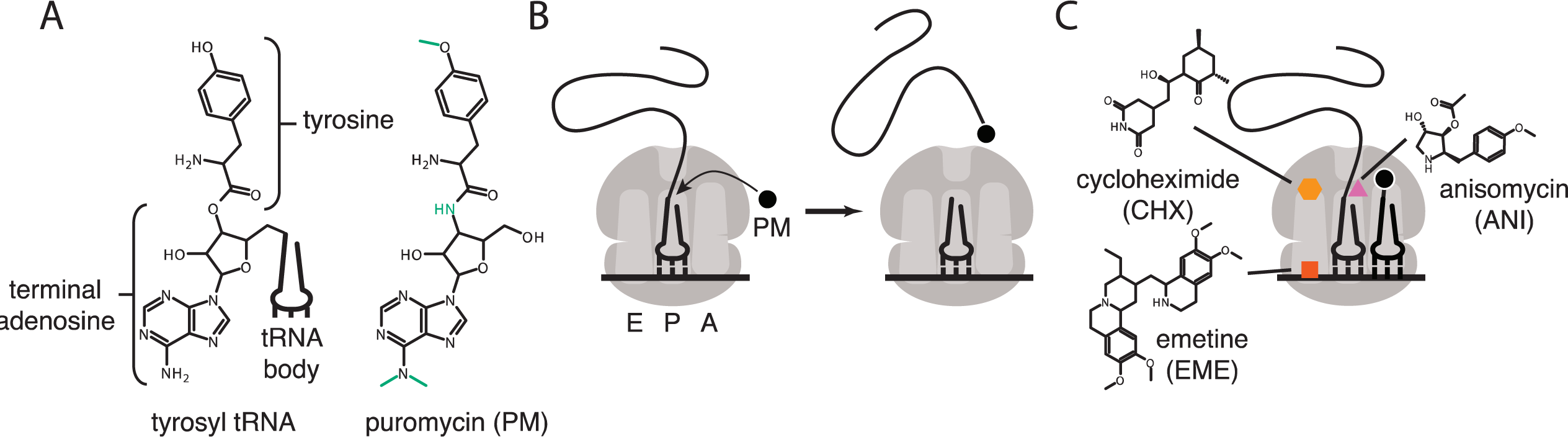
(A) Comparison of structure of 3′ terminus of tyrosyl tRNA with that of puromycin. Key differences are highlighted in green. tRNA body not drawn to scale. (B) Scheme for reaction of puromycin with peptidyl P-site tRNA on the ribosome, leading to dissociation of puromycylated peptide. (C) Structures and schematicized ribosome binding sites of translational inhibitors cycloheximide, anisomycin and emetine. Binding sites are based on *(43, 44)*.

The development of anti-puromycin antibodies and of derivatized analogs of puromycin (*4, 5*) has led to the commonplace use of puromycin as a metabolic probe to measure the extent of active translation, replacing radioactive tracers such as S35 methionine. These probes can be used to quantify the amount of active translation from cells in a culture dish, tissue or organism (*6, 7*). Subsequent development of the ribopuromycylation method (RPM) (*8*) pushed the technique a step further, claiming to detect the subcellular localization of actively translating ribosomes using a puromycin-specific antibody. In these initial publications, the authors argued that the translation elongation inhibitors cycloheximide or emetine prevent dissociation of the puromycylated peptides from the ribosome. Cycloheximide and emetine, however, bind in the E site of the ribosome (Figure 1C) far from the peptidyl-transferase center (*9, 10*). Previous work (*11*) established that these inhibitors prevent puromycin-induced splitting of ribosomes into individual subunits, but do not prevent the release of the majority of puromycylated peptides. Cycloheximide and emetine, in fact, are sometimes omitted from *in vivo* puromycylation assays based on the claim that “short” (minutes) labeling times capture peptides near their original site of translation before significant diffusion has taken place. Puromycin-based imaging methods have been widely adopted, particularly in neurobiology, where translation in neuronal processes, far from the cell body, is crucial to neuronal function (*12–20*). Other studies have combined puromycin treatment with proximity-dependent ligation (PLA) to monitor the location of translation of a specific protein (*12*), but this method again does not address diffusion of puromycylated peptides post-release from the ribosome.

In the present work, we establish that puromycin-based methods, as currently implemented, do not accurately localize translation at the subcellular level. We used a rabbit reticulocyte lysate system to show that puromycin nearly instantaneously releases nascent proteins from the ribosome, and that this release reaction is completely unaffected by emetine. To validate this finding in cells, we visualized sites of active translation using fixed cell single-molecule imaging with the SunTag reporter system. Brief treatment with puromycin nearly completely dissociated nascent peptides from their mRNAs, again, independent of the presence of emetine. Simple diffusion calculations predict that the released peptides could diffuse to nearly any point within even large mammalian cells within seconds or minutes. Thus, puromycylation methods described in the literature do not establish subcellular localization of translation.

## Results

### The puromycin analog OPP labels nuclei in live *C. elegans* germlines in the presence or absence of emetine

O-propargyl-puromycin (OPP) is a click-reactive cell permeable puromycin analog that is commonly used to localize sites of translation (*5*). When incubated with live cells or tissues, OPP reacts with translating ribosomes and becomes covalently attached to elongating peptides. Post-labeling, OPP is detected by click-reactive chemistry which attaches a fluorescent probe to OPP (Figure 2). Using this method to label translation in live *C. elegans* gonads, we observed bright labeling of live germlines upon a 5-minute incubation with OPP. The OPP signal was most intense in nuclei, specifically in the chromatin-free center where nucleoli reside. A lower signal was also observed in the cytoplasm, which contains the majority of (if not all) functional ribosomes (*21*). OPP labeling of nuclei was ablated by pre-treatment with anisomycin, a competitive inhibitor of puromycin that stops elongation by binding to the peptidyl-transferase center (*22*). In contrast, OPP labeling was unaffected by pre-treatment with emetine (Figure 2B). Emetine-resistant puromycin labeling of nucleoli has been observed previously in tissue culture cells (*8*) and may reflect trafficking or diffusion of puromycylated peptides into the nucleolus (*23, 24*). We conclude that OPP labels translational products but does not necessarily identify sites of active translation even in the presence of emetine.

**Fig. 2:**
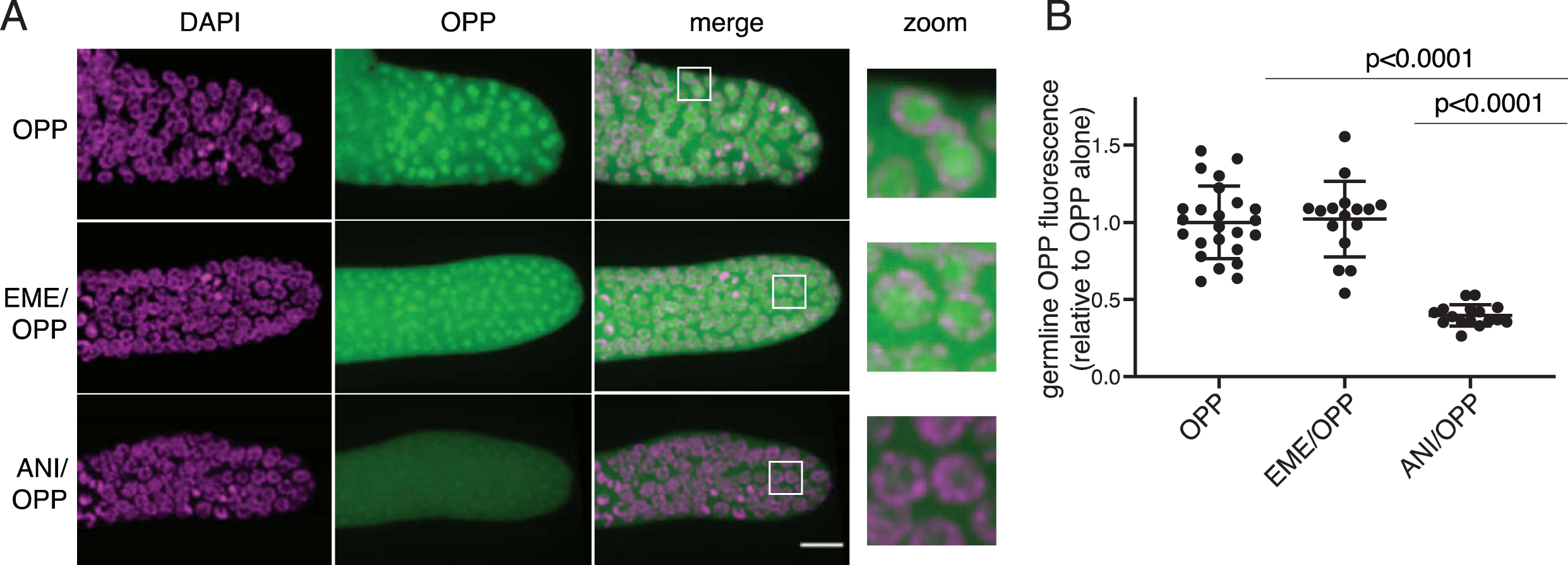
O-propargyl-puromycin (OPP) labels nuclei in the distal germline of C. elegans in the presence or absence of emetine. (A) Representative photomicrographs of germlines labeled for 5 minutes with 20 *μ*M OPP, and pre-treated for 15 minutes with control buffer (top row), 45 *μ*M emetine 15 (second row), or 37 *μ*M anisomycin (bottom row). DAPI labels chromosomes. Post-fixation, click labeling of OPP with Alexa Fluor 488 picolyl azide revealed OPP throughout the cytoplasm and concentrated in nuclei. Scale bar = 10*μ*m. (B) Quantification of OPP-Alexa 488 signal in distal germlines. Each dot represents the average fluorescence of the mitotic zone of one worm germline. Values are normalized to the average obtained for germlines pre-treated with control buffer (OPP alone). P values were obtained through an unpaired t test.

### Emetine does not prevent release of puromycylated peptides in rabbit reticulolysates

To determine whether emetine prevents release of puromycylated nascent peptides *in vitro*, we made use of a previously established real-time translation monitoring assay in rabbit reticulocyte lysate (RRL). This method relies on the fact that luciferase rapidly folds into an enzymatically active conformation, but only after release from the ribosome (*25, 26*). By programming reticulocyte lysate with a luciferase mRNA that is truncated just upstream of the stop codon, we accumulate stalled ribosomes at the 3’ end of the mRNA, in which the luciferase nascent peptide remains ribosome-bound and enzymatically inactive (Figure 3A). RRL programmed with the truncated mRNA (yellow trace) displays little luciferase activity compared to RRL translating full-length mRNA (purple trace). Addition of 91 μM puromycin (the same concentration used by (*8*)) caused a sharp increase in luminescence output from the truncated mRNA, consistent with release of the stalled peptides.

**Fig. 3:**
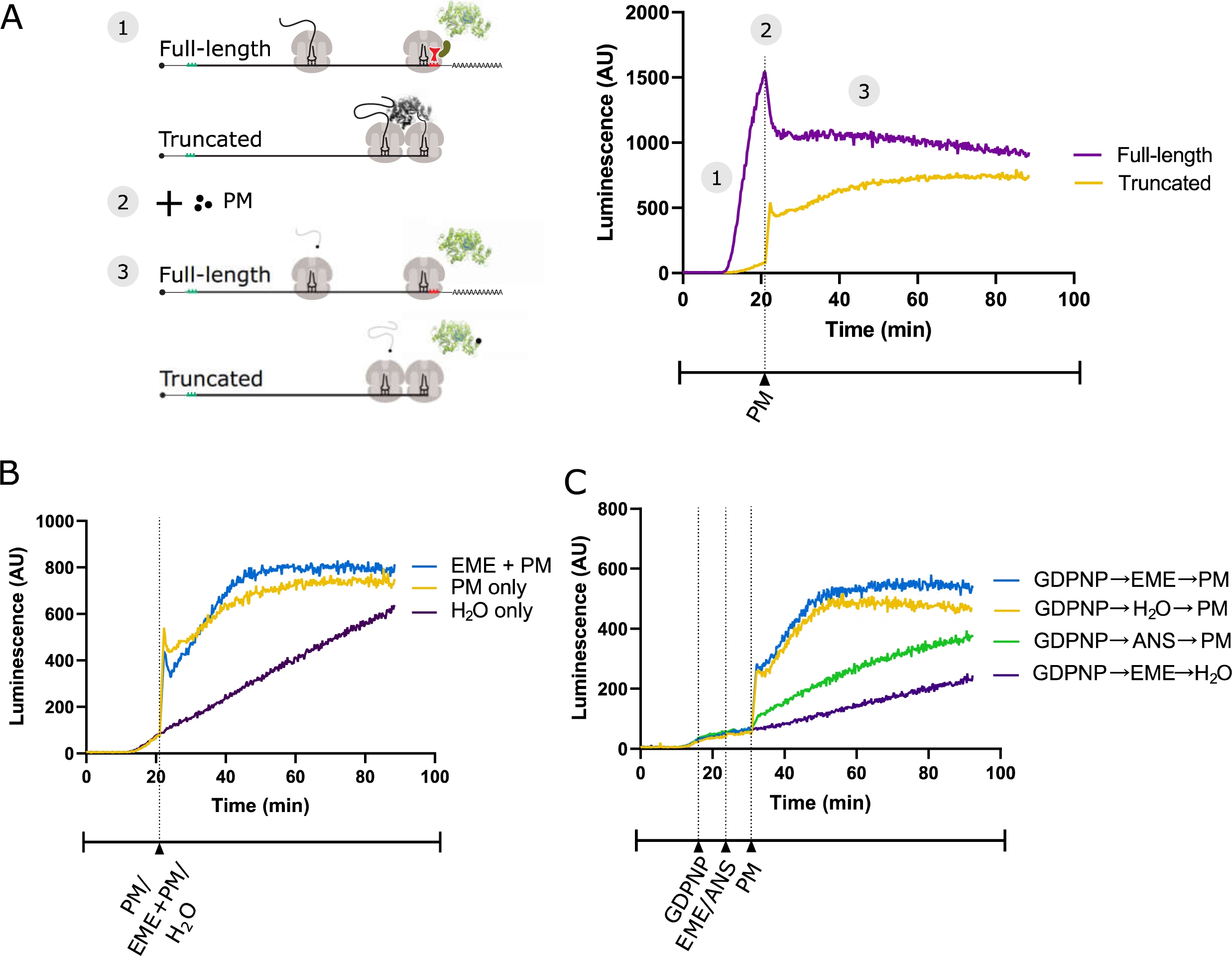
(A) Schematic of the real-time translation monitoring assay in rabbit reticulocyte lysate. (1) (Purple trace) Ribosomes translate the full-length luciferase mRNA and release luciferase which becomes enzymatically active and results in an increase in luminescence. (Yellow trace) Ribosomes stall at the 3’ end of a truncated luciferase mRNA and little to no luminescence is observed as the ribosome-bound luciferase peptides are in an enzymatically inactive conformation. (2) Puromycin (PM) is added to the system, stopping further translation and causing all nascent peptides to release from the ribosomes. (3) (Yellow) The luciferase rapidly folds into an enzymatically active conformation and a substantial increase in luminescence is observed. (B) Either puromycin (blue), H2O (purple) or a mixture of emetine (EME) and puromycin (yellow) was added to the reaction at *t* = 21 min. Experiment was performed in duplicate; mean traces are shown. (C) GDPNP was added to the reaction at *t* = 16 min for five minutes to inhibit translation across samples. Then, either emetine (blue, purple), anisomycin (ANI) (green) or H2O (yellow) is added to the reaction followed by puromycin (blue, yellow, green) or H2O (purple) five minutes later. Experiment was performed in duplicate; mean traces are shown. Note that the experiments in Figures 3A, 3B, and S1B were done in the same batch, and the yellow traces (puro treated) in these panels are the same.

We reasoned that if emetine prevents release of puromycylated nascent peptides, then emetine should block the increase in luminescence observed upon puromycin addition. Matching the conditions of David et al. (*8*), we treated the RRL with 208 μM emetine and 91 μM puromycin. When added separately, these two drugs effectively inhibit translation of full-length luciferase mRNA encoding a normal stop codon (Figure S1A)(*8*). Upon addition of puromycin, we noticed the expected steep increase in luminescence (yellow trace) that was not inhibited by simultaneous addition of emetine (blue trace) (Figure 3B). The luminescence of the no-puromycin control (purple trace) increased slowly over time, likely due to low levels of ribosome rescue activity (*27*) or spontaneous peptidyl-tRNA hydrolysis in the lysate.

We next considered the possibility that blocking peptide release with emetine requires pre-incubation. To test this, we pre-treated the lysate with emetine 5 minutes before addition of puromycin. Because pre-treatment would decrease the total translation time and overall luminescence of a sample, it was critical to equalize the total uninhibited reaction time of all samples. This was accomplished in two different ways. In a first experiment, we treated all samples with the nonhydrolyzable GTP analog 5’-guanylyl imidodiphosphate (GDPNP) to inhibit the translational GTPases and prevent ongoing translation while leaving the ribosome free to react with emetine and puromycin (Figure 3C). Again, puromycin treatment (yellow trace) caused a sharp increase in luminescence that was not affected by emetine pre-treatment (blue trace) but was inhibited by pre-treatment with anisomycin (green trace). The residual slow increase in the anisomycin trace is likely due to incomplete inhibition by anisomycin resulting from its stochastic dissociation during the reaction. In a second experiment, we added puromycin for the puromycin-only control at the same time that we started pretreating the other samples with inhibitors. This effectively inhibited translation in all samples at the same time (Figure S1B). While the increase in luminescence for the puromycin-only control (yellow trace) occurred earlier than for the pretreated samples, once the puromycin was added, the luminescence activity of the emetine pretreated sample (blue trace) matched that of the puromycin-only control. Taken together, these results show that pretreating translating ribosomes with emetine does not prevent the release of nascent peptides by puromycin *in vitro*.

### Emetine does not prevent release of puromycylated peptides in cells

To directly test whether emetine blocks release of puromycylated nascent chains *in vivo*, we implemented the SunTag method for monitoring translation on single mRNAs (*28–32*). This technique relies on a reporter mRNA encoding tandem repeats of the SunTag epitope near the 5′ end of the coding sequence (Figure 4A). When translated, each SunTag peptide is bound by a single chain variable fragment (scFV) of a GCN4 antibody fused to super folder GFP (scFV-sfGFP). An auxin-inducible degron (AID) near the 3′ end of the coding sequence allows controlled degradation of the fully-synthesized SunTag array, reducing fluorescence background and enabling detection of single fully-synthesized polypeptides. We performed fixed-cell imaging of cells stably expressing both the SunTag reporter and scFV-sfGFP, detecting mRNA by fluorescence in-situ hybridization (FISH) and SunTag signal by immunofluorescence (IF). With this single-molecule FISH and IF hybrid assay (smFISH-IF), we quantified the association of SunTag nascent chains with their encoding mRNAs under various treatment conditions.

**Fig. 4:**
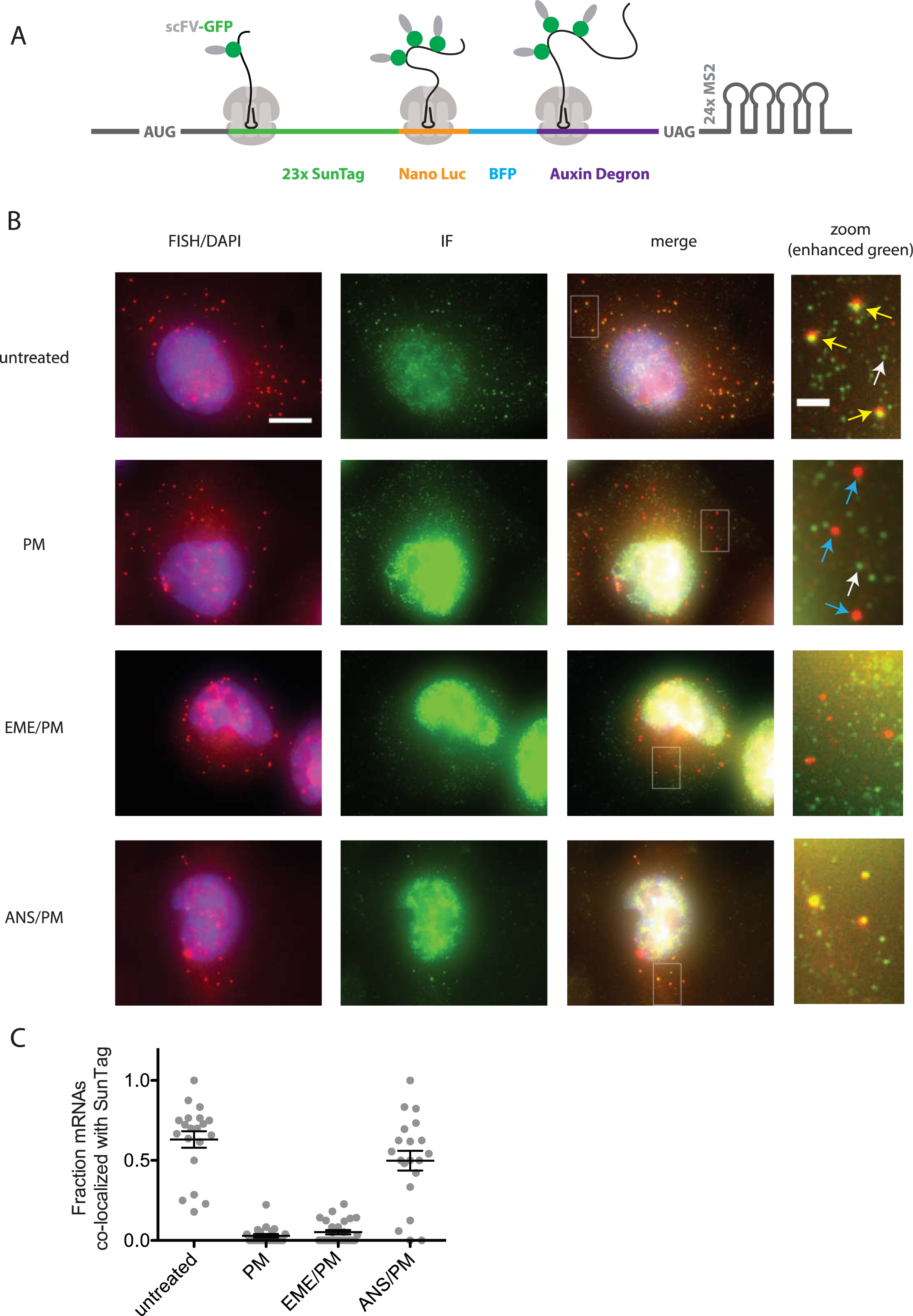
(A) SunTag reporter schematic. In addition to the tandem SunTag repeats and the auxin-inducible degron, this reporter encodes nano luciferase and BFP, which are not used in the present experiments. The 3’ UTR also encodes tandem repeats of the MS2 stem loop, which can be used to label the mRNA red. However, since we detect mRNA by FISH, we do not use the MS2 stem loops in the present experiments. (B) Example cells imaged by FISH-IF. Cells were either untreated (top row), treated with 91 μM puromycin for 5 min (second row), pre-treated with 208 μM emetine for 15 min followed by 91 μM puromycin for 5 min (third row), or pre-treated with 37 μM anisomycin for 5 min followed by 91 μM puromycin for 5 min (last row). Yellow arrows: examples of translating mRNAs; White arrows: example of single fully synthesized SunTag polypeptide (released from the ribosome); Blue arrows: examples of untranslating mRNAs. Scale bar in top left image: 10 microns. Scale bar in top right image: 2 microns. (C) Fraction of mRNAs co-localized with SunTag signal. Each dot represents one cell. Cells are only included in the analysis if they have more than 5 and fewer than 35 mRNAs. Black lines indicate mean with standard error of the mean.

In untreated cells, an average of 63% of single mRNAs (red foci) per cell co-localize with bright SunTag signal (green foci) (Figure 4B, top row and 4C); these co-localized spots reflect mRNAs bound by ribosomes synthesizing the SunTag reporter, while weaker isolated green spots reflect single fully synthesized SunTag polypeptides that have been released from the ribosome (*29*). Upon treatment with 91 μM puromycin for 5 min, an average of only 3% of mRNAs per cell colocalize with green signal, consistent with release of nascent chains upon puromycin treatment. Remarkably, pre-treatment with 208 μM emetine yielded similar results: only 5% of mRNAs on average colocalized with SunTag signal. Importantly, pre-treatment with the elongation inhibitor anisomycin, resulted in an average of 50% of mRNAs co-localized with green foci, as seen in untreated cells. Together, these data indicate that 5 minutes of puromycin treatment causes release of nascent polypeptides and diffusion away from ribosomes. Pre-treatment with emetine has no effect on puromycin-induced release.

### Puromycylation treatment times are long compared to protein diffusion rates

While initial reports argued that emetine was required to stabilize the interaction of puromycylated peptides with ribosomes, some recent studies of local protein synthesis via the puromycylation method relied on treatment with puromycin alone for ∼5-10 min, with the implication that detected nascent proteins do not appreciably diffuse away from their site of synthesis (i.e. ribosome) within the treatment time (*13, 14, 33*). To determine how far a nascent protein might diffuse on these timescales (i.e. the spatial resolution of the method), we calculated the expected displacement as a function of time, based on the previously measured diffusion coefficient of GFP in the cytosol (*34*) (Figure 5). This calculation depends on the dimensionality of space in which the molecule is confined. However, even in the most limiting case of one-dimensional diffusion—approximating movement along a very narrow neural projection—a protein is expected to diffuse ∼100 um in less than 1 min. This distance is large compared to both the scale of the relevant structures to which protein synthesis was localized in neurons (tens of microns) (*12, 13*), and to the diameter of HeLa cells (∼20 microns) (*35*), in which the method was demonstrated (*8*). Thus, limiting puromycin treatment time to a few minutes does not ensure that nascent proteins remain confined to the subcellular region in which they are synthesized.

**Fig. 5:**
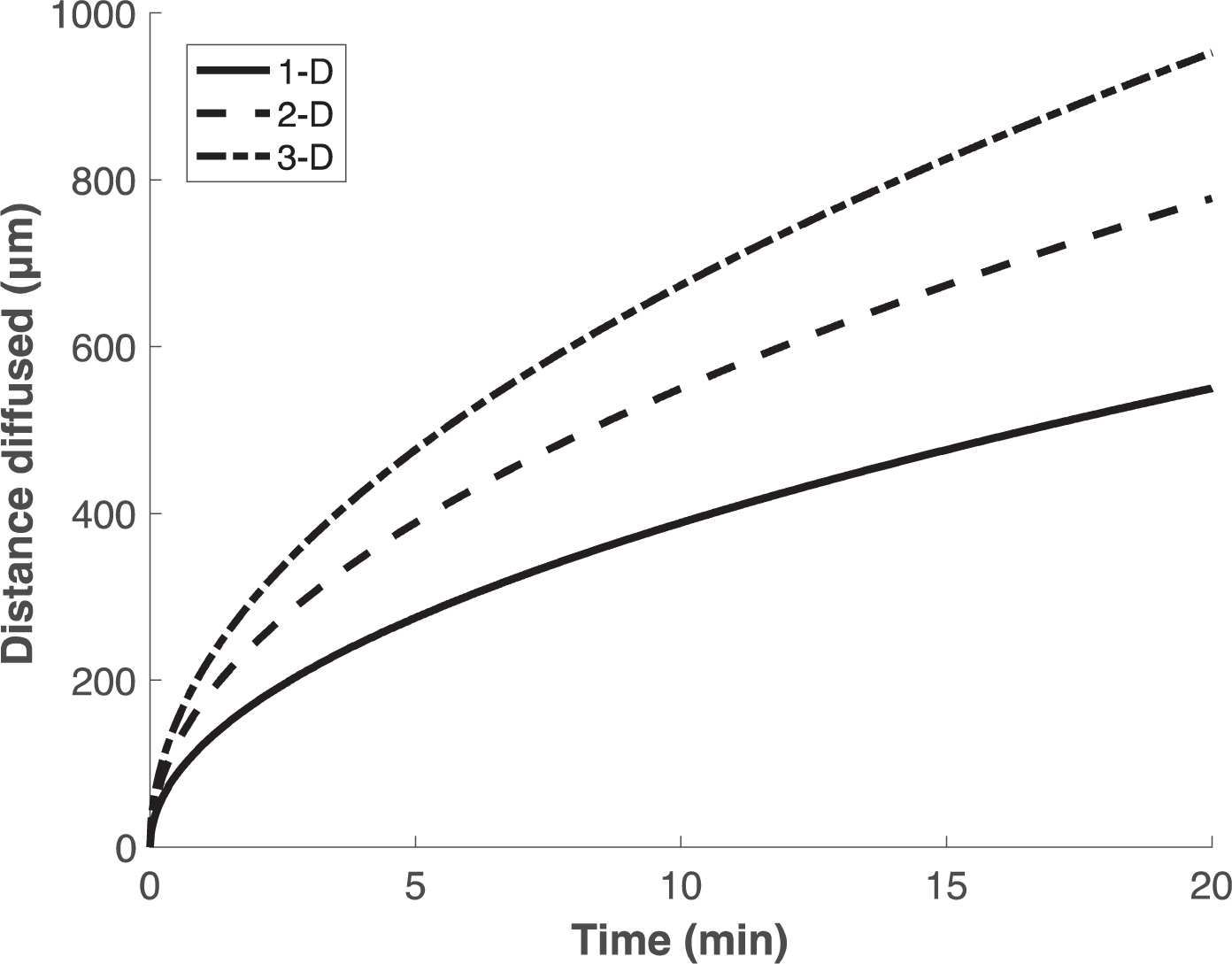
Calculation of expected displacement by diffusion as a function of time, using the equation *<x*^*2*^*> =2nDt*, where *n* is the dimensionality, *D* is the diffusion coefficient (126 um^2^/s (34)) and t is time. The calculation is shown for 1, 2 and 3 dimensions.

## Discussion

In this work, we have demonstrated that the puromycin method for visualizing localized translation does not faithfully detect nascent polypeptides at the site of their synthesis. Puromycylated polypeptides are released from the ribosome and can diffuse far away from the site of synthesis, even following short treatment times. Additionally, treatment with emetine does not prevent release of puromycylated peptides from the ribosome. It is therefore likely that the specific subcellular localizations detected by this method are in many cases the end result of trafficking or diffusion of partially-synthesized puromycylated proteins. For example, the bright nucleolar labeling detected in (*8*) likely does not reflect nucleolar translation, but the trafficking of puromycylated N-terminal fragments of highly abundant ribosomal proteins to the nucleolus (*23, 24*). Thus, conclusions reached using this method should be treated cautiously, even in neurons, where cellular projections protrude relatively far from the cell body. We note that other reporter-based methods that rely on rapid, single turnover chemistry may allow for more accurate localization of the sites of protein synthesis (*36*). Additionally, in principle, derivatizing puromycin with a chemical moiety large enough to obstruct its passage through the ribosome exit tunnel would immobilize reacted nascent chains on the ribosome. The viability of this concept is demonstrated by the RiboLace method (*37*), which uses a puromycin-biotin conjugate bound to magnetic beads to capture translating ribosomes from cellular lysates. Of course, these beads are not cell permeable and are unsuitable for *in vivo* imaging. The methods outlined here will be useful for screening cell-permeable puromycin derivatives for their ability to faithfully localize protein synthesis.

## Materials and Methods

### Plasmid construction

A Kozak consensus sequence (GCCACC) was inserted immediately upstream of the start codon of luciferase reporter plasmid pGEM-luc (GenBank X65316.2) to generate pSL312. This was used as the template for full-length firefly luciferase mRNA transcription and was linearized with a StuI restriction digest. For reasons unrelated to the current work, a disabled 2A peptide sequence was fused downstream of the luciferase sequence to generate P3.28_pGEM_ luc_2A_AGP|_kozak_RC, which was further modified by inserting an HpaI restriction site 2 nt 3′ of the final codon of luciferase (TTGtt|aac, where TTG is the final luciferase sense codon) through site-directed mutagenesis to make P3.35_pGEM_luc_trunc_kozak_RC. This plasmid was linearized with an HpaI restriction digest such that transcription of this template would terminate at TTGtt and would exclude the 2A peptide sequence.

### OPP-click

*C. elegans* was cultured according to standard methods at 20°C. N2 adult germlines were dissected into egg buffer with 1mM levamisole. Germlines were incubated with 45 *μ*M emetine, or 37 *μ*M anisomycin (*38*), or egg buffer alone for 15 minutes. OPP was added at a concentration of 20 *μ*M while maintaining concentrations of emetine and anisomycin for the 5 minute incubation. Germlines were rinsed once with PBS and fixed in 4% paraformaldehyde. Click reaction was carried out with Click-iT Plus OPP Alexa Fluor 488 Protein Synthesis Assay kit (Thermo Fisher C10456) according to the manufacturer’s directions.

### C. elegans Imaging

Images were taken with a Zeiss Axio Observer equipped with a CSU-W1 SoRA spinning disk scan head (Yokogawa) and Slidebook v6.0 software (Intelligent Imaging Innovations). Germline images are 10 *μ*m z-stacks starting at the bottom of the distal germline with 0.27 *μ*m step size using a 63X objective. Images were acquired with laser power set to 20, and exposure time set to 25 ms in the 488 nm channel and 50 ms in the 405 nm channel. Average intensity projections were quantified in ImageJ. An ROI was drawn around the mitotic zone of each germline and fluorescence in the 488 nm channel was measured. Fluorescence intensity of each germline was normalized to the average intensity of the germlines treated with OPP alone.

### Luciferase-based real-time translation monitoring assay

Luciferase plasmids were linearized with a blunt-end restriction enzyme just upstream (truncated) or downstream (full-length) of the stop codon, followed transcription with the mMESSAGE mMACHINE SP6 transcription kit (Invitrogen AM1344). Synthesized mRNA was quantified using a Nanodrop 1000. Nuclease-treated rabbit reticulocyte lysate translation reactions (Promega L4960) were set up in a 384-well plate (Thermo Scientific 164610) on ice. Luciferin (PerkinElmer 122799) was added to each reaction well to a concentration of 0.5mM followed by 12 units of Superase-In RNase Inhibitor (Invitrogen AM2696). SP6-transcribed truncated or full-length firefly luciferase mRNA was added to a concentration of 40ug/mL using a multichannel pipette and the plate was immediately inserted into a luminometer microplate reader (Biotek Synergy H1MD) regulated at 30°C. Luminescence readings were taken every few seconds, depending on the number of reaction wells. 5’-guanylyl imidodiphosphate (GDPNP; Jena Bioscience NU-401-50) was added to the wells 16 minutes after the start of the reaction for five minutes at a concentration of 100uM followed by a five-minute pretreatment of either 208uM emetine (Cayman Chemical 21048) or 9.4uM anisomycin (Sigma A9789). Puromycin (Sigma Aldrich P7255) was added to wells at a concentration of 91uM. In experiments where GDPNP was not used, the first translation inhibitors were added to the reaction wells at 21 minutes following the start of the reaction.

### smFISH Probe Labeling

smFISH probes targeting the SunTag region of the mRNA reporter transcript were synthesized as described(*39, 40*). 20-mer olgionucleotides were ordered from IDT in an arrayed format, pooled, and labeled on the 3’-end with amino-11-12 ddUTP (Lumiprobe A5040) using deoxynucleotidyl transferase (TdT, Thermo Fisher EP0162). After size exclusion purification on a Spin-X centrifuge column (Corning 8161) with Bio Gel P-4 Beads (Bio Rad 1504124), the oligonucleotide was labeled with Cy3. Following the labeling reaction, the probes were again purified over a Spin-X column to remove excessive dyes.

### smFISH-IF

smFISH-IF was performed similarly as described(*39, 41*). smFISH-IF was performed on cell stably expressing the SunTag mRNA reporter and scFV-sfGFP. 18 mm #1 coverslips (Fisher 12-545-100) were etched in 3M sodium hydroxide (Millepore Sigma 221465) prior to cell plating. The coverslips were then washed 3x with PBS (Corning 21-031-CV) and then coated for 30 min at 37°C with 0.25 mg/mL rat tail collagen I (Gibco A1048301) diluted in 20 mM sodium acetate (Sigma-Aldrich S2889). After another 2x PBS wash, 18,000 cells were plated per well and grown for 24 hours in DMEM supplemented with 10% FBS. 24 hours following plating, the media was supplemented with 1 ug/mL doxycycline hyclate (Millipore Sigma, #D9891) and 500 uM 3-indole acetic acid (IAA) (Sigma-Aldrich I2886).

Approximately 24 hours following induction, cells were treated with either 91 uM puromycin in the media for 5 min, 208 uM emetine in the media for 15 min followed by 91 uM puromycin in the media for 5 min, or 37 uM anisomycin in the media for 5 min followed by 91 uM puromycin in the media for 5 min. Control cells were left untreated. Following treatment, samples were prepared for smFISH-IF. All solutions were prepared in nuclease free water (Quality Biological 351-029-131CS). Cells were washed 3x with 1x PBS (Corning 46-013-CM) + 5 mM magnesium chloride (Sigma-Aldrich M2670-500G) (PBSM). Cell were then fixed for 10 minutes at room temperature in PBSM + 4% paraformaldehyde (Electron Microscopy Sciences 50-980-492). Following fixation, samples were washed for 3×5 minutes in PBSM and permeabilized for 10 minutes in PBSM + 5 mg/mL BSA (VWR VWRV0332-25G) + 0.1% Triton-X100 (Sigma-Aldrich T8787-100mL). After 3×5 minute washes in PBSM, cell were incubated for 30 minutes at room temperature in 2xSSC (Corning 46-020-CM), 10% formamide (Sigma-Aldrich F9037-100ML), and 5 mg/mL BSA (VWR VWRV0332-25G). Following pre-hybridization incubation, samples were incubated for 3 hours at 37°C in 2xSSC (Corning 46-020-CM), 10% formamide (Sigma-Aldrich F9037-100ML), 1 mg/mL competitor E. coli tRNA (Sigma-Aldrich 10109541001), 10% w/v dextran sulfate (Sigma-Aldrich D8906-100G), 2 mM ribonucleoside vanadyl complex (NEB S1402S), 100 units/mL SUPERase In (Thermo Fisher AM2694), 60 nM SunTag_v4-Cy3 smFISH probes, and 1:1,000 chicken anti-GFP (Aves Labs GFP-1010). The coverslips were then washed 4x with 2xSSC (Corning 46-020-CM) + 10% formamide (Sigma-Aldrich F9037-100ML). The samples were then incubated with 2×20 minutes with a goat anti-chicken IgY secondary antibody labeled with Alexa Fluor 488 (Thermo Fisher A-11039). After 3x washes in 2xSCC, cells were mounted on pre-cleaned frosted glass cover slides (Fisher 12-552-3) with ProLong Diamond antifade reagent with DAPI (Invitrogen P36962). After curing for 24 hours, the samples were imaged on a custom Nikon Ti-2 wide-field microscope equipped with a 60x 1.4NA oil immersion objective lens (Nikon), a Spectra X LED light engine (Lumencor), and a Orca 4.0 v2 scMOS camera (Hamamatsu). The microscope was under automated control by Nikon Elements software. x-y pixel size: 107.5 nm. z-step: 300 nm.

### smFISH Analysis

Fixed cell image analysis was performed as described(*39*) with existing or custom MATLAB software. Spot detection of the mRNA and proteins channels were performed independently in FISH-Quant(*42*). In the protein channel, all released single peptides in the cytoplasm were detected and thresholded based on their gaussian fitting parameters (intensity and width) and inspected to ensure accuracy. All released single peptides were then averaged into an idealized point spread function to calculate the integrated intensity of a single SunTag array. In the mRNA channel, only cytoplasmic RNAs were included for analysis. After determining all cytoplasmic mRNA positions, FISH-Quant’s transcription site quantification algorithm was employed to quantify the integrated intensity of the associated translation site. Briefly, a 11×11 bounding box was drawn at the position of each mRNA and Gaussian fitting was performed centered on the brightest pixel within this box. The integrated intensity of the translation site was then normalized against the intensity of the idealized single peptide to calculate the number of nascent chains associated with a given mRNA. The translation sites were filtered based on shape, intensity, and distance from the mRNA. Failure to converge on an accurate fit given these parameters resulted in the associated translation site intensity to have an intensity value of 0. Translation sites with an integrated intensity of less than one idealized single peptide were determined to be unassociated with SunTag signal. Only cells with greater than 5 and fewer than 35 mRNAs were considered.

## Acknowledgements

This work was funded by the National Institutes of Health [2R37GM059425-14 to R.G.; 5R37HD037047-20 to G.S.; 5K99GM135450-02 to B.Z] and the Howard Hughes Medical Institute (HHMI) (R.G. and G.S.). B.Z. was an HHMI fellow of the Damon Runyon Cancer Research Foundation [DRG-2250-16] for a portion of this study. D.H.G. is a Damon Runyon Fellow supported by the Damon Runyon Cancer Research Foundation (DRG-2280-16). N.M.L. and M.C. are supported by NIH Training Grant T32 GM007445. M.C. is supported by National Science Foundation Graduate Research Fellowship DGE-1746891.

## Supplemental Figures

**Fig. S1:**
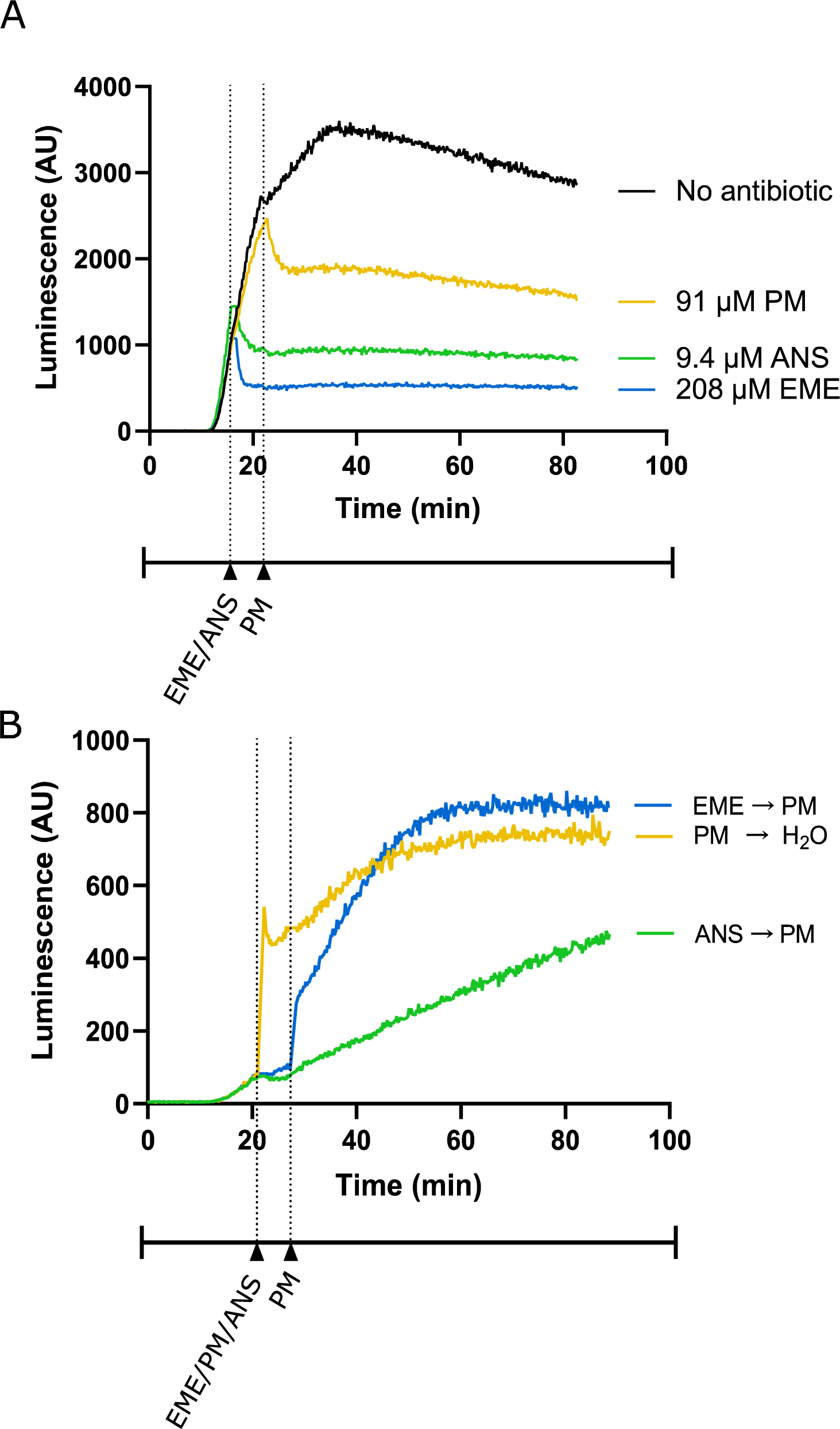
(A) Luminescence measured from translation of full-length luciferase mRNA in rabbit reticulocyte lysate. Due to Either emetine (blue), anisomycin (green) or puromycin (yellow) was added to the reaction at indicated concentrations. experimental restrictions, emetine and anisomycin were added at *t* = 16 min and puromycin was added at *t* = 22 min. (B) Luminescence measured from translation of truncated luciferase mRNA in rabbit reticulocyte lysate. Either emetine (blue), puromycin (yellow) or anisomycin (green) was added to the reaction at *t* = 21 min for five minutes to inhibit translation across the samples. Then, either puromycin (blue, green) or H2O (yellow) is added to the reaction. Experiment was performed in duplicate; mean traces are shown.

## References

1. R. Aviner, The science of puromycin: From studies of ribosome function to applications in biotechnology. Comput. Struct. Biotechnol. J. 18 (2020), pp. 1074–1083.

2. M. B. Yarmolinsky, G. L. D. L. Haba, Inhibition by puromycin of amino acid incorporation into protein. Proc. Natl. Acad. Sci. 45, 1721–1729 (1959).

3. D. Nathans, Puromycin inhibition of protein synthesis: incorporation of puromycin. Proc. Natl. Acad. Sci. United States. 51, 585–592 (1964).

4. K. Fujiwara A. Ogawa, H. Asada, H. Saikusa, H. Nakamura, S. Ono, T. Kitagawa, The Preparation of Puromycin Antibody and Its Use in Enzyme Immunoassay for the Quantification Using β-D-Galactosidase as a Label. J. Biochem. 92, 1599–1605 (1982).

5. J. Liu, Y. Xu, D. Stoleru, A. Salic, Imaging protein synthesis in cells and tissues with an alkyne analog of puromycin. Proc. Natl. Acad. Sci. U. S. A. 109, 413–418 (2012).

6. E. K. Schmidt, G. Clavarino, M. Ceppi, P. Pierre, SUnSET, a nonradioactive method to monitor protein synthesis. Nat. Methods. 6, 275–277 (2009).

7. J. A. Chao, Y. J. Yoon, R. H. Singer, Imaging translation in single cells using fluorescent microscopy. Cold Spring Harb. Perspect. Biol. 4 (2012), doi: 10.1101/cshperspect.a012310.

8. A. David, B. P. Dolan, H. D. Hickman, J. J. Knowlton, G. Clavarino, P. Pierre, J. R. Bennink, J. W. Yewdell, Nuclear translation visualized by ribosome-bound nascent chain puromycylation. J. Cell Biol. 197, 45–57 (2012).

9. N. Garreau De Loubresse, I. Prokhorova, W. Holtkamp, M. V. Rodnina, G. Yusupova, M. Yusupov, Structural basis for the inhibition of the eukaryotic ribosome. Nature. 513, 517–522 (2014).

10. W. Wong, X. C. Bai, A. Brown, I. S. Fernandez, E. Hanssen, M. Condron, Y. H. Tan, J. Baum, S. H. W. Scheres, Cryo-EM structure of the Plasmodium falciparum 80S ribosome bound to the anti-protozoan drug emetine. Elife. 2014 (2014), doi: 10.7554/eLife.03080.

11. A. P. Grollman, Inhibitors of protein biosynthesis. V. Effects of emetine on protein and nucleic acid biosynthesis in HeLa cells. J. Biol. Chem. 243, 4089–94 (1968).

12. S. Tom Dieck, L. Kochen, C. Hanus, M. Heumüller, I. Bartnik, B. Nassim-Assir, K. Merk, T. Mosler, S. Garg, S. Bunse, D. A. Tirrell, E. M. Schuman, Direct visualization of newly synthesized target proteins in situ. Nat. Methods. 12, 411–414 (2015).

13. A. Biever, C. Glock, G. Tushev, E. Ciirdaeva, T. Dalmay, J. D. Langer, E. M. Schuman, Monosomes actively translate synaptic mRNAs in neuronal processes. Science (80-.). 367 (2020), doi: 10.1126/science.aay4991.

14. C. Glock, A. Biever, G. Tushev, I. Bartnik, B. Nassim-Assir, S. tom Dieck, E. M. Schuman, bioRxiv, in press, doi: 10.1101/2020.06.09.141960.

15. P. V’kovski, M. Gerber, J. Kelly, S. Pfaender, N. Ebert, S. Braga Lagache, C. Simillion, J. Portmann, H. Stalder, V. Gaschen, R. Bruggmann, M. H. Stoffel, M. Heller, R. Dijkman, V. Thiel, Determination of host proteins composing the microenvironment of coronavirus replicase complexes by proximity-labeling. Elife. 8 (2019), doi: 10.7554/eLife.42037.

16. Y. E. Lewis, A. Moskovitz, M. Mutlak, J. Heineke, L. H. Caspi, I. Kehat, Localization of transcripts, translation, and degradation for spatiotemporal sarcomere maintenance. J. Mol. Cell. Cardiol. 116, 16–28 (2018).

17. J. J. Langille, K. Ginzberg, W. S. Sossin, Polysomes identified by live imaging of nascent peptides are stalled in hippocampal and cortical neurites. Learn. Mem. 26, 351–362 (2019).

18. T. Gonatopoulos-Pournatzis, R. Niibori, E. W. Salter, R. J. Weatheritt, B. Tsang, S. Farhangmehr, X. Liang, U. Braunschweig, J. Roth, S. Zhang, T. Henderson, E. Sharma, M. Quesnel-Vallières, J. Permanyer, S. Maier, J. Georgiou, M. Irimia, N. Sonenberg, J. D. Forman-Kay, A. C. Gingras, G. L. Collingridge, M. A. Woodin, S. P. Cordes, B. J. Blencowe, Autism-Misregulated eIF4G Microexons Control Synaptic Translation and Higher Order Cognitive Functions. Mol. Cell. 77, 1176-1192.e16 (2020).

19. T. E. Graber, S. Hébert-Seropian, A. Khoutorsky, A. David, J. W. Yewdell, J. C. Lacaille, W. S. Sossin, Reactivation of stalled polyribosomes in synaptic plasticity. Proc. Natl. Acad. Sci. U. S. A. 110, 16205–16210 (2013).

20. T. E. Graber, E. Freemantle, M. N. Anadolu, S. Hébert-Seropian, R. L. Macadam, U. Shin, H. D. Hoang, T. Alain, J. C. Lacaille, W. S. Sossin, UPF1 governs synaptic plasticity through association with a STAU2 RNA granule. J. Neurosci. 37, 9116–9131 (2017).

21. S. Klinge, J. L. Woolford, Ribosome assembly coming into focus. Nat. Rev. Mol. Cell Biol. 20 (2019), pp. 116–131.

22. A. P. Grollman, Inhibitors of protein biosynthesis. II. Mode of action of anisomycin. J. Biol. Chem. 242, 3226–33 (1967).

23. S. Kubota, T. D. Copeland, R. J. Pomerantz, Nuclear and nucleolar targeting of human ribosomal protein S25: Common features shared with HIV-1 regulatory proteins. Oncogene. 18, 1503–1514 (1999).

24. C. Schmidt, E. Lipsius, J. Kruppa, “Nuclear and Nucleolar Targeting of Human Ribosomal Protein S6” (1995).

25. V. A. Kolb, E. V Makeyev, A. S. Spirin, Folding of firefly luciferase during translation in a cell-free system. EMBO J. 13, 3631–7 (1994).

26. J. Frydman, E. Nimmesgern, K. Ohtsuka, F. U. Hartl, Folding of nascent polypeptide chains in a high molecular mass assembly with molecular chaperones. Nature. 370, 111–7 (1994).

27. S. Shao, K. von der Malsburg, R. S. Hegde, Listerin-Dependent Nascent Protein Ubiquitination Relies on Ribosome Subunit Dissociation. Mol. Cell. 50, 637–648 (2013).

28. X. Pichon, A. Bastide, A. Safieddine, R. Chouaib, A. Samacoits, E. Basyuk, M. Peter, F. Mueller, E. Bertrand, Visualization of single endogenous polysomes reveals the dynamics of translation in live human cells. J. Cell Biol. 214, 769–81 (2016).

29. B. Wu, C. Eliscovich, Y. J. Yoon, R. H. Singer, Translation dynamics of single mRNAs in live cells and neurons_Wu_Singer_2016. 352 (2016).

30. T. Morisaki, K. Lyon, K. F. DeLuca, J. G. DeLuca, B. P. English, Z. Zhang, L. D. Lavis, J. B. Grimm, S. Viswanathan, L. L. Looger, T. Lionnet, T. J. Stasevich, Real-time quantification of single RNA translation dynamics in living cells. Science (80-.). 352, 1425–1429 (2016).

31. C. Wang, B. Han, R. Zhou, X. Zhuang, Real-Time Imaging of Translation on Single mRNA Transcripts in Live Cells. Cell. 165, 990–1001 (2016).

32. X. Yan, T. A. Hoek, R. D. Vale, M. E. Tanenbaum, Dynamics of Translation of Single mRNA Molecules in Vivo. Cell. 165, 976–989 (2016).

33. C. E. Holt, K. C. Martin, E. M. Schuman, Local translation in neurons: visualization and function. Nat. Struct. Mol. Biol. 26 (2019), pp. 557–566.

34. C. Di Rienzo, V. Piazza, E. Gratton, F. Beltram, F. Cardarelli, Probing short-range protein Brownian motion in the cytoplasm of living cells. Nat. Commun. 5, 1–8 (2014).

35. A. B. Borle, Kinetic analyses of calcium movements in HeLa cell cultures. I. Calcium influx. J. Gen. Physiol. 53, 43–56 (1969).

36. Y. Na, S. Park, C. Lee, D. K. Kim, J. M. Park, S. Sockanathan, R. L. Huganir, P. F. Worley, Real-Time Imaging Reveals Properties of Glutamate-Induced Arc/Arg 3.1 Translation in Neuronal Dendrites. Neuron. 91, 561–573 (2016).

37. M. Clamer, T. Tebaldi, F. Lauria, P. Bernabò, R. F. Gómez-Biagi, M. Marchioretto, D. T. Kandala, L. Minati, E. Perenthaler, D. Gubert, L. Pasquardini, G. Guella, E. J. N. Groen, T. H. Gillingwater, A. Quattrone, G. Viero, Active Ribosome Profiling with RiboLace. Cell Rep. 25, 1097-1108.e5 (2018).

38. A. Bastide, J. Yewdell, A. David, The RiboPuromycylation Method (RPM): an Immunofluorescence Technique to Map Translation Sites at the Sub-cellular Level. BIO-PROTOCOL (2018), doi: 10.21769/bioprotoc.2669.

39. D. H. Goldman, N. M. Livingston, J. Movsik, B. Wu, R. Green, bioRxiv, in press, doi: 10.1101/2020.05.29.121954.

40. I. Gaspar, F. Wippich, A. Ephrussi, Enzymatic production of single-molecule FISH and RNA capture probes. RNA. 23, 1582–1591 (2017).

41. M. J. Latallo, N. M. Livingston, B. Wu, Translation imaging of single mRNAs in established cell lines and primary cultured neurons. Methods. 162–163, 12–22 (2019).

42. F. Mueller, A. Senecal, K. Tantale, H. Marie-Nelly, N. Ly, O. Collin, E. Basyuk, E. Bertrand, X. Darzacq, C. Zimmer, FISH-quant: automatic counting of transcripts in 3D FISH images. Nat. Methods. 10, 277–278 (2013).

43. N. Garreau De Loubresse, I. Prokhorova, W. Holtkamp, M. V. Rodnina, G. Yusupova, M. Yusupov, Structural basis for the inhibition of the eukaryotic ribosome. Nature. 513, 517–522 (2014).

44. W. Wong, X. C. Bai, A. Brown, I. S. Fernandez, E. Hanssen, M. Condron, Y. H. Tan, J. Baum, S. H. W. Scheres, Cryo-EM structure of the Plasmodium falciparum 80S ribosome bound to the anti-protozoan drug emetine. Elife. 2014 (2014), doi: 10.7554/eLife.03080.

